# Top-down feedback matters: Functional impact of brainlike connectivity motifs on audiovisual integration

**DOI:** 10.1101/2024.10.01.615270

**Authors:** Mashbayar Tugsbayar, Mingze Li, Eilif B. Muller, Blake Richards

**Affiliations:** McGill University, Montréal, Canada; Department of Neurosciences, Faculty of Medicine, Université de Montréal, Montréal, Canada; Centre de Recherche Azrieli du CHU Sainte-Justine, Montréal, Canada; Mila Quebec AI Institute, Montréal, Canada

## Abstract

Artificial neural networks (ANNs) are an important tool for studying neural computation, but many features of the brain are not captured by standard ANN architectures. One notable missing feature in most ANN models is top-down feedback, i.e. projections from higher-order layers to lower-order layers in the network. Top-down feedback is ubiquitous in the brain, and it has a unique modulatory impact on activity in neocortical pyramidal neurons. However, we still do not understand its computational role. Here we develop a deep neural network model that captures the core functional properties of top-down feedback in the neocortex, allowing us to construct hierarchical recurrent ANN models that more closely reflect the architecture of the brain. We use this to explore the impact of different hierarchical recurrent architectures on an audiovisual integration task. We find that certain hierarchies, namely those that mimic the architecture of the human brain, impart ANN models with a light visual bias similar to that seen in humans. This bias does not impair performance on the audiovisual tasks. The results further suggest that different configurations of top-down feedback make otherwise identically connected models functionally distinct from each other, and from traditional feedforward and laterally recurrent models. Altogether our findings demonstrate that modulatory top-down feedback is a computationally relevant feature of biological brains, and that incorporating it into ANNs affects their behavior and constrains the solutions it’s likely to discover.

## 1 Introduction

Artificial neural networks (ANNs) often draw inspiration from the brain and serve in turn as tools with which to model it (Doerig et al., 2023). For example, deep convolutional neural networks (CNN)–originally inspired by hierarchical feature detection in the mammalian brain (Fukushima, 1980)—predict behavior and representations in visual cortex with high accuracy (Cadena et al., 2019; Yamins & DiCarlo, 2016). However, there are still significant gaps in most deep neural networks’ ability to capture behavior, particularly when presented with ambiguous, challenging stimuli (Geirhos et al., 2018; Kar et al., 2019). This gap is thought to be a result, in part, of the exclusively feedforward structure of CNNs, as opposed to the biological brain, which consists of feedforward as well as local and top-down recurrent connections (Kar et al., 2019; van Bergen & Kriegeskorte, 2020). Local recurrence is modeled in a subfamily of ANNs known as recurrent neural networks (RNNs), and there have been studies using RNNs to model sensory processes in the brain (Kubilius et al., 2018). In contrast, top-down feedback has largely been neglected in deep models, with a few exceptions (Islah et al., 2023; Naumann et al., 2022; Tsai et al., 2024; Wybo et al., 2022). It’s unclear whether omitting top-down feedback changes the behavior of a model or has an impact on processing in a deep model.

As such, we built ANNs with biologically-inspired top-down feedback and examined whether different configurations of feedback vs feedforward connectivity had any relevant effect on model behavior and preferences. Importantly, top-down feedback connections are functionally and physiologically distinct from feedforward connections. They typically connect higher order association areas to lower order sensory areas (Felleman & Van Essen, 1991). Data suggests that top-down inputs in the neocortex are modulatory (Larkum, 2004; Sherman & Guillery, 1998), meaning they alter the magnitude of neural activity, but do not drive it themselves. It would be beneficial for computational neuroscience if ANN models allowed us to explore the potential unique impacts of top-down feedback on computation in the neocortex.

For this study, we chose to examine the effects of ANNs with top-down feedback on an audiovisual categorization task. We chose audiovisual tasks because feedback connections can also span sensory modalities, and the auditory areas (which project directly to V1 in primates (Clavagnier et al., 2004)) are speculated to sit higher on the sensory-association hierarchy than primary visual areas (King & Nelken, 2009). As such, audio modulation of visual processing likely occurs via top-down feedback across multiple cortical sites. Audiovisual interplay produces many well-known functional effects, such as the McGurk effect (McGurk & Macdonald, 1976) and sound-induced flash illusions (Shams et al., 2000), making it a particularly interesting testbed for studies of the computational role of top-down feedback.

To study the effect of top-down feedback on such tasks, we built a freely available code base for creating deep neural networks with an algorithmic approximation of top-down feedback. Specifically, top-down feedback was designed to modulate ongoing activity in recurrent, convolutional neural networks. We explored different architectural configurations of connectivity, including a configuration based on the human brain, where all visual areas send feedforward inputs to, and receive top-down feedback from, the auditory areas. The human brain-based model performed well on all audiovisual tasks, but displayed a unique and persistent visual bias compared to models with only driving connectivity and models with different hierarchies. This qualitatively matches the reported visual bias of humans engaged in audio-visual tasks (Posner et al., 1976; Stokes & Biggs, 2014). Our results confirm that distinct configurations of feedforward/feedback connectivity have an important functional impact on a model’s behavior. Therefore, top-down feedback captures behaviors and perceptual preferences that do not manifest reliably in feedforward-only networks. Further experiments are needed to clarify whether top-down feedback helps an ANN fit better to neural data, but the results show that top-down feedback affects the processing of stimuli and is thus a relevant feature that should be considered for deep ANN models in computational neuroscience more broadly.

## 2 Results

### 2.1 Framework for modeling top-down feedback with RNNs

Based on neurophysiological data, we implemented a deep neural network model where each layer corresponds to a brain region and receives two different types of input: feedforward and feedback. Feedforward input drives the activity of the layer as it does in regular ANNs. In contrast, inspired by the physiology of apical dendrites in pyramidal neurons, feedback input to each neuron is integrated and then run through another non-linear function. The result of this is then multiplied with the integrated feedforward activation (Fig. 1A, left). As such, feedback input cannot activate a neuron that is not receiving feedforward input, nor can it decrease the activity of the neuron, but it can modulate its level of activity (Fig. 1A, right). This mimics the modulatory role of top-down feedback in the neocortex as observed in neurophysiological experiments (Larkum, 2004; McAdams & Maunsell, 1999). But, we note that experimental and modeling research suggests that top-down feedback can sometimes be weakly driving (Larkum, 2004; Reynolds et al., 2000; Shai et al., 2015), i.e. it can reduce the threshold for activation for the neuron, a form of feedback referred to as “composite” feedback (Shai et al., 2015). We explore the impact of both purely multiplicative feedback and composite feedback below.

**Figure 1:**
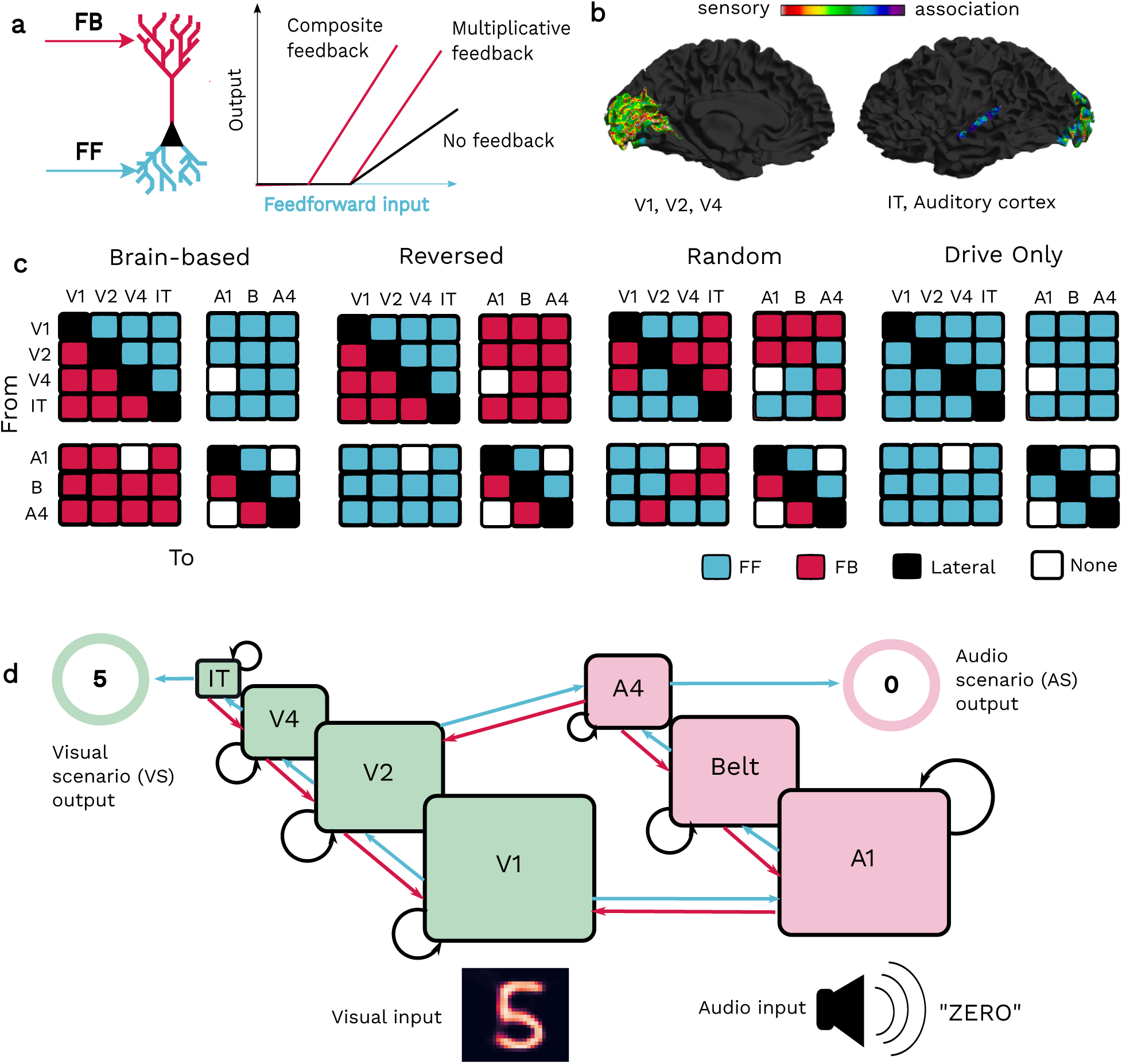
Description of models. **a**, Each area of the model receives driving feedforward input and modulatory feedback input. Feedback input alters the gain of a neuron, but it doesn’t affect its threshold of activation (multiplicative feedback). In later experiments, we explore an alternative mechanism, where it weakly affects the threshold of activation (composite feedback). **b**, Modeled regions and their externopyra- midisation values (i.e thickness and relative differentiation of supragranular layers, used as proxy measure for sensory-associational hierarchical position). Note higher overall externopyramidisation values in the occipital lobe compared to the temporal lobe. **c**, Using above hierarchical measures, we constructed models where each connection has a direction (i.e regions send either feedforward or feedback connections to other regions). In the brainlike model based on human cytoarchitectural data, where all visual regions send feedforward to and receive feedback from the auditory regions. In the reverse model, all auditory regions send feedforward to and receive feedback from visual regions, while connections within a modality remain the same. **d**, The resulting ANN. Outputs of image identification tasks are read out taken from IT, outputs of audio identification tasks are read out from A4, an auditory associational area. Connections between modules are simplified for illustration.

We implemented retinotopic and tonotopic organization, as well as local recurrence, by using a Convolutional Gated Recurrent Unit (GRU) to model each brain region (Ballas et al., 2016; Cho et al., 2014). These models were end-to-end differentiable, and therefore, we could train them with backpropagation-of-error. Importantly, there is nothing about this modeling framework that demands a specific type of training, meaning that the models created within this framework can undergo supervised, unsupervised, or reinforcement learning, though here we used supervised tasks (see Methods).

For the audiovisual categorization tasks, we constructed a series of ANNs with four layers corresponding to ventral visual regions (V1, V2, V4, and IT) and three layers corresponding to ventral auditory regions (A1, Belt, A4). For one ANN, which we will call the “brainlike” model, we used the externopyramidisation of each cortical area as seen in human histological data to determine the hierarchical position of said area in the network (Fig. 1B). Externopyramidisation refers to the relative differentiation of the supragranular layers in the cortex (Sanides, 1962). Percentage of supragranular projecting neurons (SLN) is a longstanding measure of hierarchical distance in mammals, based on experimental observations that long-range feedforward connections originate primarily from the supragranular layers, while feedback connection are infragranular in origin (Barone et al., 2000; Felleman & Van Essen, 1991; Markov & Kennedy, 2013). Externopyramidisation is an indirect estimate of supragranular projecting neurons, based on observations in primates that feedforward-projecting sensory areas feature dense, highly differentiated supragranular layers (Gerbella et al., 2007) while feedback-projecting transmodal areas have less differentiated supragranular layers and larger infragranular layers (Morecraft et al., 2012). Researchers thus assume that higher externopyramidisation scores (i.e. thicker and more differentiated supragranular layers) indicates a “lower order“ region that sends more feedforward connections (Goulas et al., 2018). We make the same assumption in-line with previous literature quantifying hierarchical distance from human histological data (Paquola et al., 2019; Saberi et al., 2023), but we note that our modeling framework could easily use other metrics for determining position in the cortical hierarchy (Paquola et al., 2020; Wagstyl et al., 2015; Zilles & Amunts, 2009).

This resulted in an architecture wherein visual areas provided feedforward input to all auditory areas, whereas auditory areas provided feedback to visual areas (Fig. 1C, left), similar to an existing hypothesis on primate audiovisual processing (King & Nelken, 2009). In another ANN, that we call the “reversed” model, the directionality was reversed, i.e. all auditory areas feed forward to the visual areas whereas visual areas provide feedback to auditory areas. In this model, directionality of connections within a modality were kept the same as the brain-based model (Fig. 1C, middle). In three additional control ANNs, the directionality of connections between each pair of regions was determined by a coin toss (Fig. 1C, right).

Altogether, the framework we developed allowed us to explore the functional implications of top-down feedback in multi-layer RNNs. As well, this framework allowed us to examine the functional impact of a RNN architecture inspired by the human brain on audiovisual processing, and compare it to other non-brainlike architectures.

### 2.2 Effect of top-down feedback on visual tasks with auditory cues

In order to examine the functional impact of different top-down feedback architectures, we first trained multiplicative feedback models (i.e. not composite feedback) on an image categorization task with additional auditory stimuli. The task for the ANN was to correctly identify the category based on the image, but the images were sometimes ambiguous (which were created using a variational autoencoder, see Methods). In cases where the visual image was ambiguous, the network was provided with a disambiguating auditory cue (we refer to this as Visual-dominant Stimulus case 1, or VS1; Fig. 2a, top). In addition, we presented the model with situations where the visual input was unambiguous and the auditory input was misleading (VS2; Fig. 2a, bottom), in order to test whether the models could learn to ignore misleading audio inputs. Crucially, the models were never told which stimulus scenario they were encountering. As such, they had to learn to rely on the auditory inputs when the visual inputs were ambiguous (VS1), and ignore the auditory inputs when the visual inputs were unambiguous (VS2). Models were trained in mini-batches on both VS1 and VS2 stimuli, then tested on held out data (see Methods for details).

**Figure 2:**
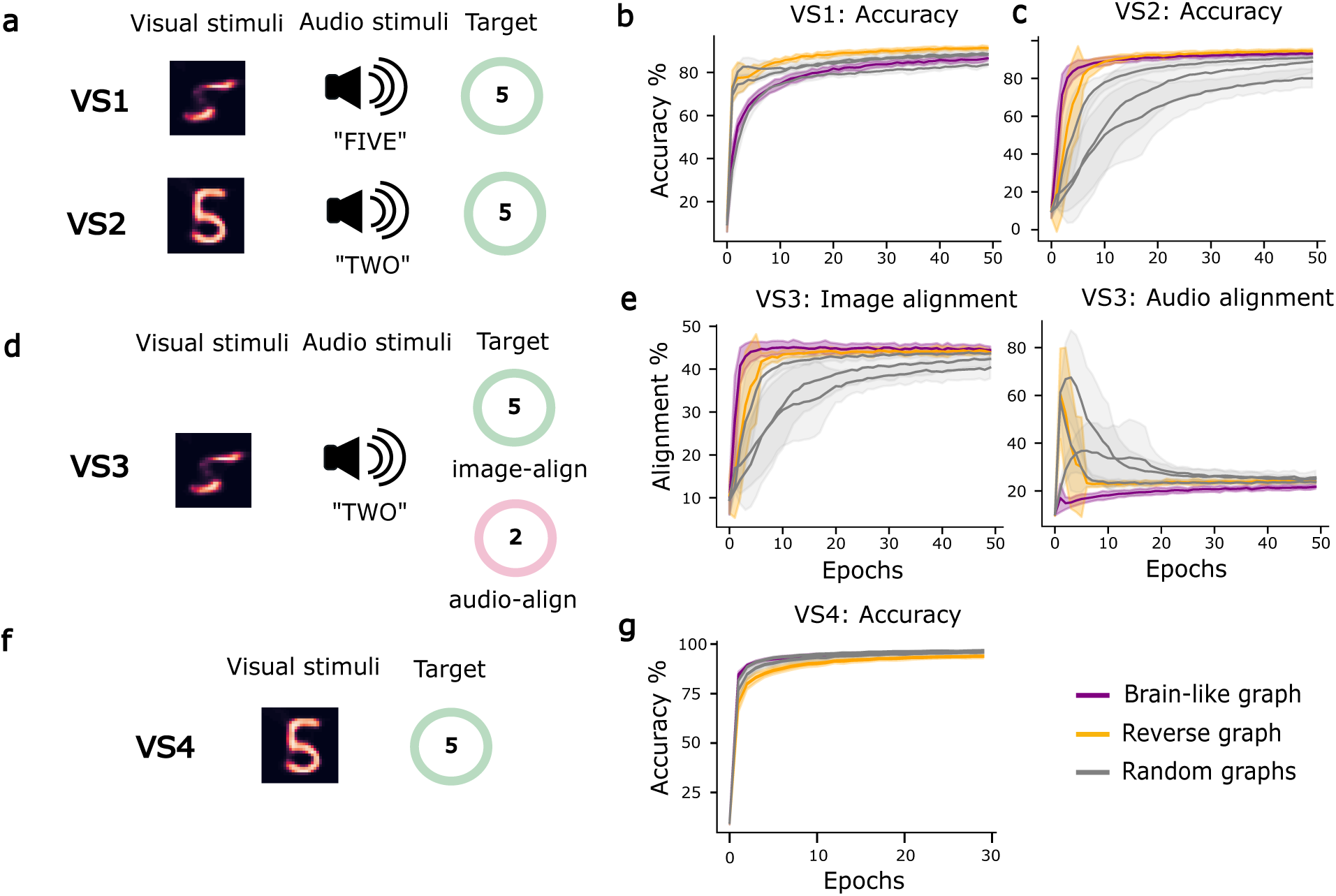
Multimodal visual tasks. **a**, Training conditions. Models must identify the visual stimulus given an ambiguous image and a matching audio clue (VS1) or unambiguous image and distracting audio (VS2). **b-c**, Accuracy across epochs for tasks VS1 and VS2 on holdout datasets, **d** Trained models were given an ambiguous visual stimulus and a nonmatching audio stimulus (VS3) to assess which modality they align most closely with. **e**, Alignment of trained models across epochs based on task VS3. Scenario was shown to model at the end of each training epoch, but never during training. **f**, Models were additionally trained and tested on image stimuli only to assess their baseline performance (VS4). **g**, Accuracy of models across epochs on task VS4

In stimulus condition VS1, all of the models were able to learn to use the auditory clues to disambiguate the images (Fig. 2b). However, the brainlike model learned to rely on auditory inputs more slowly, whereas the reversed model was exceptionally good at integrating auditory information. In comparison, in VS2, we found that the brainlike model learned to ignore distracting audio inputs quickly and consistently compared to the random models, and a bit more rapidly than the auditory information (Fig. 2c). These data show that the brainlike model is biased towards visual stimuli compared to the other models.

We next wanted to probe this visual bias further by examining how the models relied on either visual or auditory inputs when neither was unambiguously relevant to the task. Thus, in another set of test stimuli, we examined what happened when the models were presented with non-matching auditory input and ambiguous visual input (VS3; Fig. 2d). Notably, VS3 was not given as a task during training; the models had been trained on stimuli from the VS1 and VS2 conditions. In this situation, we were interested in whether the models were more likely to align their answers to the visual or auditory stimuli, given that neither provided a clear and correct answer. We found that the brainlike model aligned with the visual input from the start, while all other models had to learn to do so further along in training (Fig. 2e). Importantly, if we trained the models purely on unambiguous images (VS4; Fig. 2f), all of the models were equally adept at learning (Fig. 2g). As such, the visual bias shown by the brainlike model did not reflect a more general increased capability with visual inputs, rather, it reflected an inductive bias in audio-visual integration created by the top-down feedback architecture. Altogether, our results demonstrated that both the architecture of top-down feedback can impact audio-visual processing, and that the specific architecture inspired by the human brain has a visual bias in these tasks.

### 2.3 Effect of top-down feedback on auditory tasks with visual cues

To see if the visual bias persisted for a non-visual task, we then trained the models from scratch on an audio recognition task, where the ANNs now had to identify ambiguous sounds using visual stimuli as clues (AS1; Fig. 3a, top) and ignore distracting images when they’re not needed (AS2; Fig. 3a, bottom). As before, the models were trained on data from the AS1 and AS2 conditions, then tested on held out data points. Notably, only the brainlike model and one of the random models learned to use the visual stimuli at all, across 10 random seeds (Fig. 3b). In contrast, all of the models learned to ignore the distractor image (Fig. 3c). This indicates that the brainlike model is more inclined to use visual stimuli for the task than the other models.

**Figure 3:**
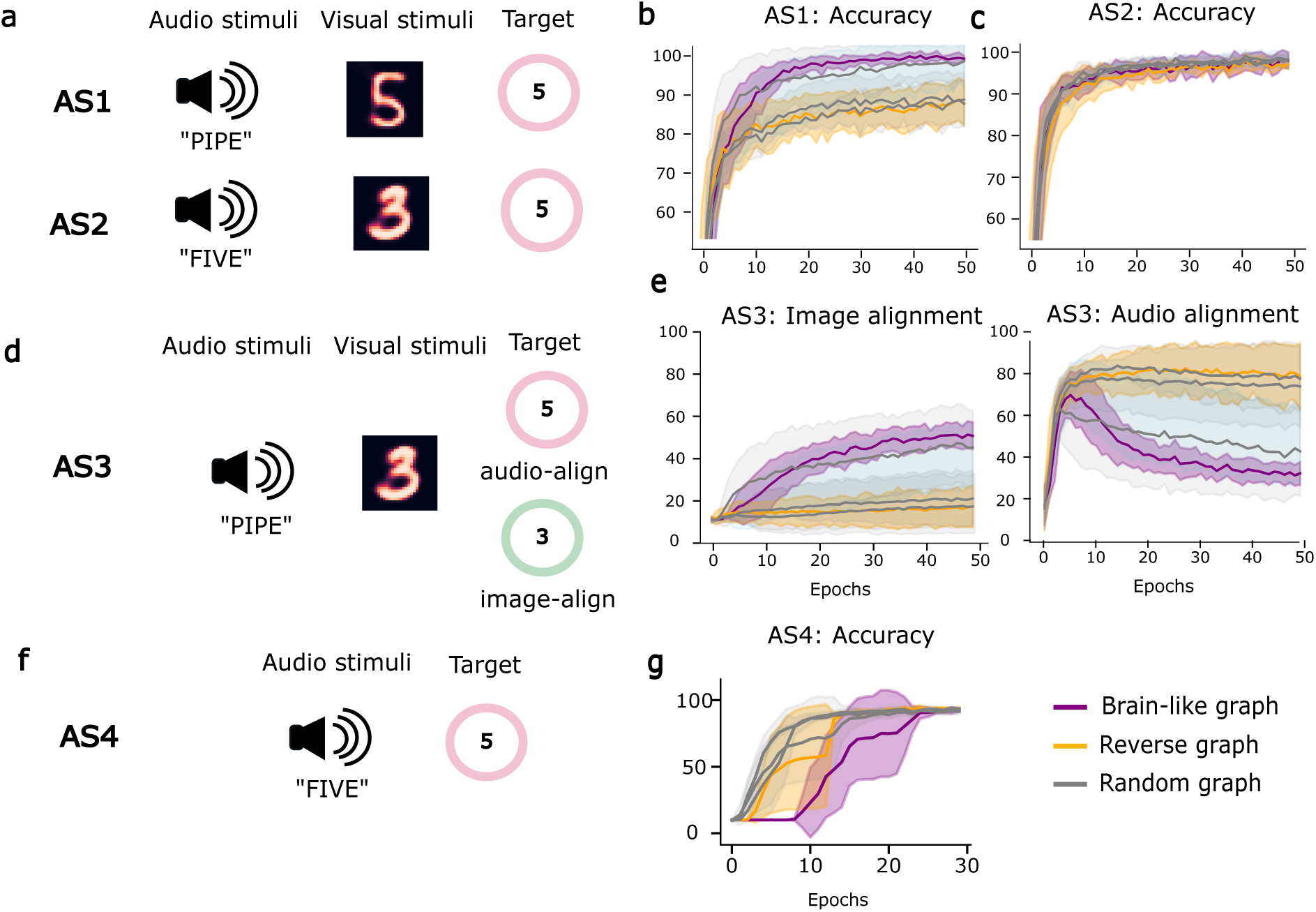
Multimodal auditory tasks. **a**, Training conditions. Models must identify the auditory stimulus given an ambiguous audio and a matching visual clue (AS1) or unambiguous audio and distracting image (AS2). **b-c**, Accuracy across epochs for tasks AS1 and AS2 on holdout datasets, **d**, Trained models were given an ambiguous audio stimulus and a nonmatching visual stimulus (AS3) to assess which modality they align most heavily with. **e**, Alignment of trained models across epochs based on task AS3. Scenario was shown to model at the end of each training epoch, but never during training. **f**, Models were additionally trained and tested on audio stimuli only to assess their baseline performance (AS4), **g**, Accuracy of models across epochs on task AS4

Next, we examined how the models responded when neither input was unambiguously informative. In a test scenario with non-matching visual stimuli and ambiguous audio (AS3; Fig. 3d), the brainlike model aligned more heavily with the visual input than most other models (Fig. 3e). Thus, despite the task being primarily auditory, the brainlike model retained its clear visual bias. Interestingly, unlike in the case of visual tasks, there was a difference between the models in terms of their ability to learn purely from unambiguous auditory stimuli (AS4; Fig. 3f). While all models could achieve the same final error rate, the brainlike model took longer to learn from purely auditory inputs than the other models. Our results suggest that the architecture of top-down feedback in the human brain may provide an inductive bias that favours visual inputs over auditory inputs.

### 2.4 Effects of composite top-down feedback

As noted above, neurophysiological evidence suggests that in addition to modulating activity in a multiplicative manner, top-down feedback in the neocortex can be weakly driving as well, shifting not just the gain but also the threshold of activation for a neuron (Fig. 1A) (Shai et al., 2015). As such, we performed the same experiments as above, but now with composite feedback, i.e. feedback that has both a multiplicative and an additive effect. Notably, composite feedback is more akin to a feedforward connection (due to the weakly driving additive component). Therefore, in order to isolate for the impact of composite top-down feedback we added a control of an identically connected network consisting only of driving feedforward connections (i.e. no top-down feedback anywhere), which we called the drive-only model. We then compared the brainlike composite feedback models to the original controls and the new drive-only control.

Interestingly, we found that composite feedback improved the performance of all models on tasks that they previously had difficulty with, while maintaining performance on the tasks they were already strong on (Fig. 4a-d). Specifically, in the visual task with ambiguous visual stimuli and informative audio cues (VS1), we observed faster learning and massively improved final accuracy in the brainlike and random models, which lagged behind the reverse model when using multiplicative feedback (Fig. 4a). Conversely, the reverse model with multiplicative feedback had difficulty using visual clues when identifying ambiguous audio (AS1). Composite feedback improved the performance of the reverse model on this task, but the already high-performing brainlike model remained unaffected by the change (Fig. 4b). The effect may be due to the slightly increased size of the composite models, as we observed that bigger models also receive a similar boost in AS1 performance (Fig. S2a). However, the composite models are 1K parameters larger than the models with multiplicative feedback (compared to an overall 280K) while the models in Fig. S2 are roughly three times the size.

**Figure 4:**
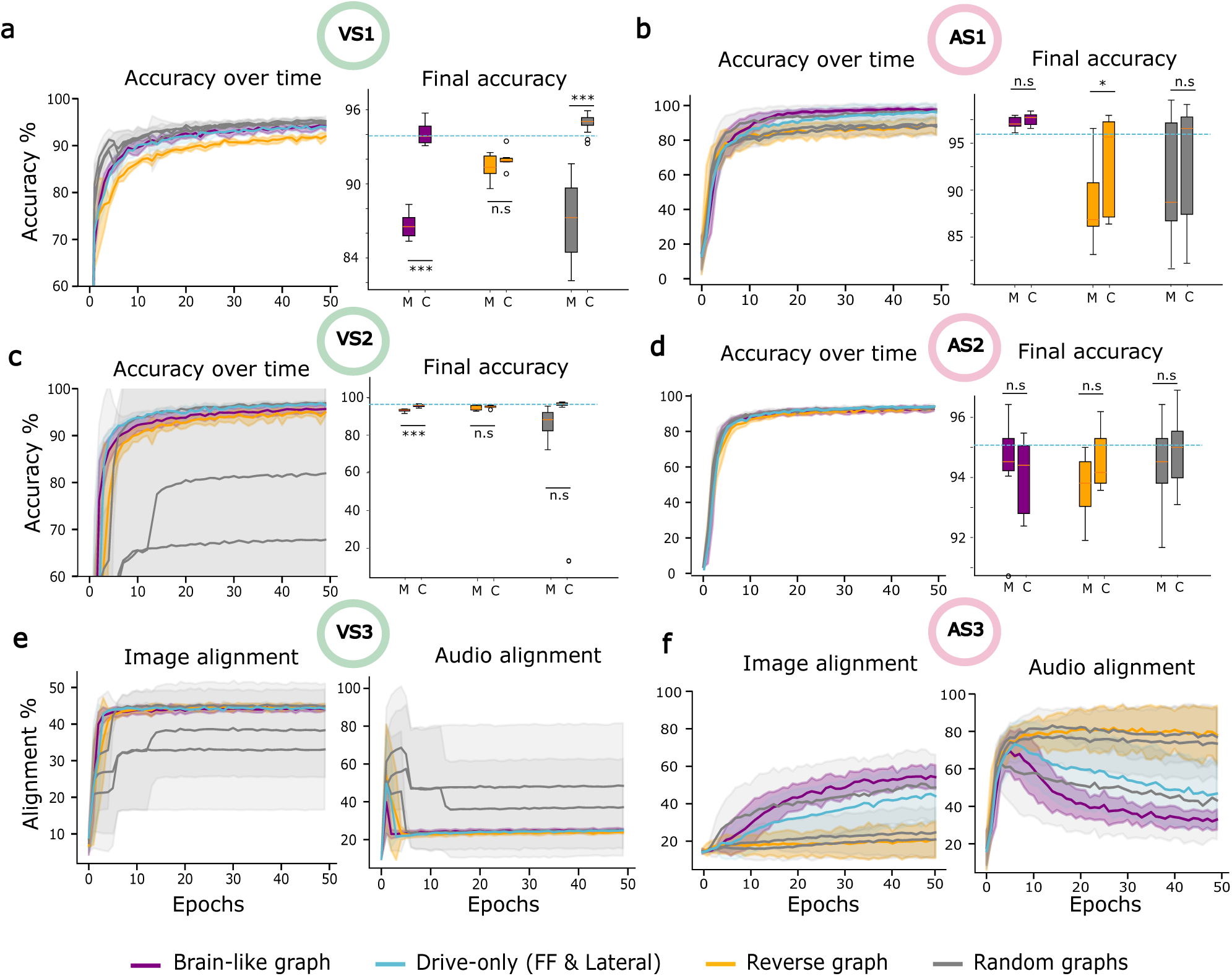
Composite versus multiplicative feedback in multimodal tasks. **a, c, e**, Test performances of models with composite feedback and drive-only models trained on visual tasks (VS1 and VS2). The final epoch accuracy of models with composite feedback (C) is compared to that of models with multiplicative feedback (M) shown in previous figures **b, d, f**, Test performances of models trained on auditory tasks (AS1 and AS2).

The impact of composite feedback was more mixed in the situation where the visual input was unambiguous and the auditory input was misleading (VS2). The brainlike model and most random models showed better final accuracy (Fig. 4c), but a few random models only performed well in VS1 and not VS2 (Fig. 4c, rightmost outliers). The only task where composite feedback had no noticeable effect was when the auditory input was unambiguous and the visual clue was misleading (AS2), likely because all models already performed at a high level with multiplicative feedback (Fig. 4d). Interestingly, drive-only models (i.e feedforward-only interlayer connections and lateral recurrence) consistently performed at an accuracy roughly equivalent to that of the brainlike composite feedback model (Fig. 4 a-d, blue lines). These data imply that composite feedback may be a more general purpose computational mechanism that can be used for efficient multi-modal learning with the correct architecture, but based on accuracy alone, it did not behave noticeably differently from a drive-only model.

We next wondered whether the difference between composite top-down feedback and the drive-only model could be observed in terms of the visual bias of the networks. Therefore, we examined the visual biases of the models in cases where no unambiguous correct input was given (VS3 and AS3). We found that in both the visual tasks (Fig. 4e) and the auditory tasks (Fig. 4f) the visual bias of the brainlike model was still present. In contrast, the drive-only model showed biases between that of brainlike and reverse models with a larger variability between seeds than the brainlike model, suggesting the drive-only model privileged the visual and auditory input based on task demands and initialization. This shows that the inductive bias towards vision in the brainlike model depended on the presence of the multiplicative component of the feedback, and therefore, the visual bias is a function of the network architecture. This visual bias shows under all conditions for the network while the bias of the drive-only model varies based on task and initialization. Altogether, these results suggest that composite feedback as seen in the neocortex is an effective mechanism for multi-modal integration, and that the architecture of feedback in the human brain provides an inductive bias towards visual stimuli.

### 2.5 Effects of top-down feedback on task-switching

Our results so far showed that top-down feedback imparts a persistent inductive bias depending on the network architecture. However, top-down feedback is thought to be particularly important for context-dependent task switching (Li et al., 2004; Liu et al., 2021). Thus, we wanted to know if the brainlike model’s visual bias lent it any advantage over other networks, particularly in tasks that required flexible responses to context.

As such, we trained the models on all four training tasks (VS1, VS2, AS1, AS2) at the same time. To do so, we augmented all the models with a new output region that takes input from IT and A4. This new region in the model can be thought of as broadly representing higher order multimodal regions in the brain (Fig. 5a). This multimodal region additionally received a binary attention flag with each pair of inputs indicating the target stimulus, i.e. whether to attend to the visual or auditory streams (Fig. 5b). As in previous tasks, the models had to determine on their own whether the extraneous stimulus was useful for identifying the target stimulus (VS1, AS1) or a distraction to be ignored (VS2, AS2).

**Figure 5:**
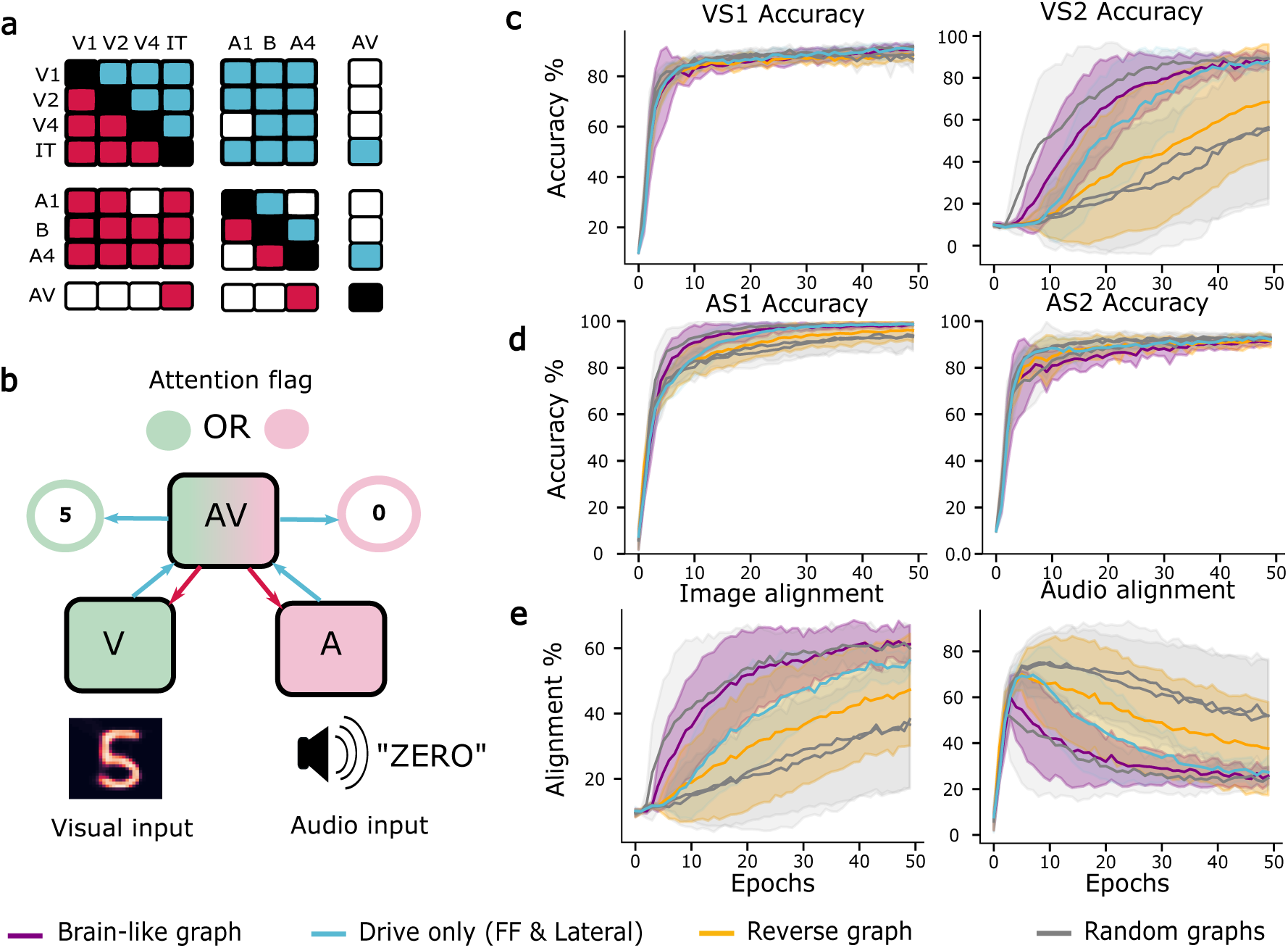
Audiovisual switching task. **a**, All models were given a new audiovisual output area (AV) connecting to IT and A4, **b**, The output area receives an attention flag telling which stream of information to attend to (visual or auditory). The models with feedback use with composite feedback. **c-d**, Test performance of models with composite feedback and drive-only models on all tasks. The models were trained simultaneously on all tasks, **e**, Alignment of models given ambiguous visual and ambiguous audio input with differing labels.

All models quickly learned to identify ambiguous visual stimuli using the auditory input (Fig. 5c, left), but the brainlike model learned to ignore distracting audio quicker than all other models including the drive-only model (Fig. 5c, right). Similarly, the brainlike model quickly learned to use visual clues when identifying ambiguous audio (Fig. 5d, left). However, it lagged behind the other models when it had to ignore the visual stimuli (Fig. 5d, right). When presented with two ambiguous stimuli of different labels, the brainlike model again aligned more frequently with the visual stimuli (Fig. 5e). In contrast, the drive-only model started out more heavily aligned to the auditory stimuli and slowly aligned with the visual stimuli through training, suggesting it conforms to task demands and does not possess the same inductive bias for visual tasks (Fig. 5e). Interestingly, the reverse model and two random models struggled to learn to ignore auditory stimuli (Fig. 5b). This is likely because the auditory dataset is smaller, less variable and thus easier to learn than the visual dataset, causing certain models to shortcut their training and rely excessively on the audio inputs (see also Fig. 3b and Fig. 3e). This data shows that a visual bias helps models ignore this shortcut, whereas an auditory bias makes them more susceptible to it. The detrimental auditory bias is suppressed during the visual only tasks (Fig. 2e), but when forced to switch between both auditory and visual tasks, the auditory bias was more apparent.

Our results show that the architectural biases imparted by feedback are even easier to observe in tasks that require flexible switching, and that the brainlike model and its visual bias could be advantageous over drive-only models in certain scenarios, if for example there is a data imbalance. In contrast, an auditory bias hampered the models for the scenarios studied here.

### 2.6 Functional specializations of model regions

Different regions of the model are active at different time steps in our simulations, depending on where the region sits in the hierarchy and when feedforward input is received by that region (Fig. 6a). We therefore wanted to understand the impact of top-down feedback on the temporal dynamics of computation in the model across different regions. As such, we next studied the representations of stimuli across time in different regions of the model, in order to gain insight into how the various regions contribute to the tasks studied here. First, to gain a qualitative understanding of representations over time in the models, we projected the hidden states of all regions of the models with composite feedback at different time points onto two dimensions using t-distributed Stochastic Neighbor Embedding (Fig. 6b, left). In general, clustered, easily separable representations greatly aid the performance of a neural network in classification tasks like these. We found that different regions showed different clustering of stimuli over time, for example, in the task where the network must classify auditory stimuli and ignore distracting audio stimuli (VS2), the one seed of the brainlike model showed a clear clustering of different stimuli in the IT region, but not in A4, and only moderately in V1 (Fig. 6b, left). To quantify the clustering pattern observed across models, we calculated the Neighborhood Hit (NH) score of all datapoints in the latent space at each time step in all regions of the model (Paulovich et al., 2008; Rauber et al., 2017).

**Figure 6:**
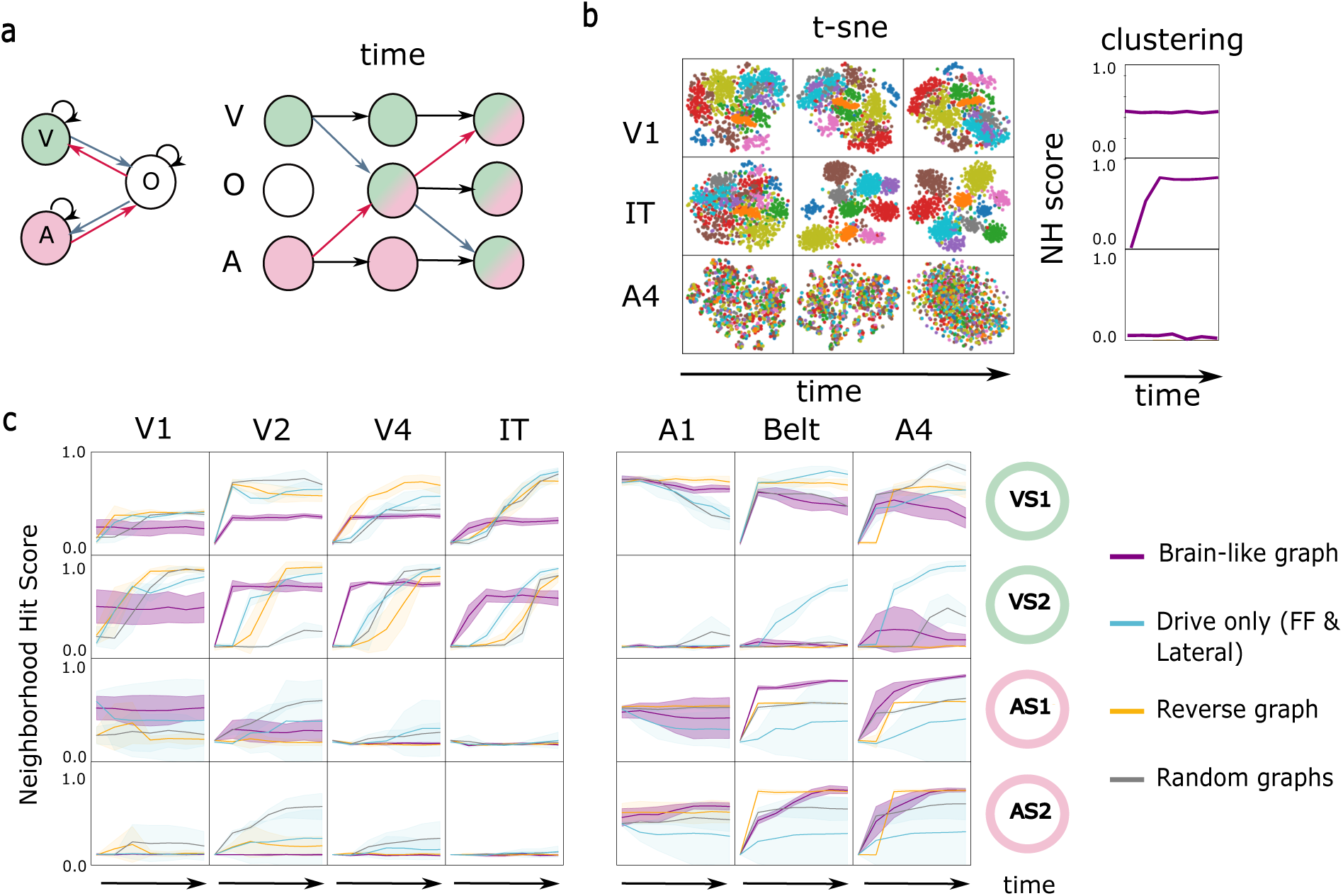
Model activity during multimodal tasks. **a**, Information flows through the model from area to area across time. At the first time step, only the primary visual and auditory areas will process information. The areas they feed forward to are activated at the next time step, incorporating top-down information if there is any. **b**, Comparison of t-SNE reduced latent space and clustering metric in three areas of the brainlike model at different time stages on task VS2 (ignore audio stimulus). **c**, Neighborhood Hit scores in all areas of the trained models across time. Trained models were taken from experiments in Fig 4

For the visual tasks, in the brainlike and reverse models, we observed distinct, clustered representations in the auditory regions when the auditory stimuli were useful for image identification (Fig. 6c, VS1). Conversely, none of the auditory regions in the brainlike and reverse models formed clustered representations in the scenario when the auditory stimuli were irrelevant to the task, outside brief clustering in the A4 in some brainlike models (Fig. 6c, VS2). Interestingly, this was not the case with the drive-only model, which exhibited clustering in all auditory regions outside A1 even when the auditory inputs were irrelevant. This demonstrates that in the models with top-down feedback there is a specialization that occurs, such that the auditory regions only clustered the data when necessary, whereas the absence of top-down feedback mechanisms led to more multimodal responses in general.

A similar delineation of function was again evident in the auditory tasks, where the brainlike model had highly clustered representations in V1 and V2 when the visual clue was useful to the task (Fig. 6c, AS1), but showed uniformly lower clustering in the visual regions after the first time step if the visual clue had to be ignored (Fig. 6c, AS2). Notably, the reverse model did not cluster the visual clue effectively resulting in low clustering in the visual regions during AS1 and a drop in performance (Fig. 3b). In contrast, clustering exhibited by the drive-only model was more variable compared to all models with top-down feedback.

It’s important to emphasize that no specific inductive biases were given to any of the regions of the model beyond the inputs they received and whether those inputs were feedforward or feedback. V2 and the auditory belt, for instance, are identical aside from their connectivity in these models. Moreover, all regions were trained end-to-end with the same, unitary loss function. As such, there was no incentive for a non-primary region to develop an exclusive visual or auditory specialization in the absence of the connectivity constraints. An especially illustrative example of this is one of the random models where the V2 region actually developed into an auditory region (Fig. 6c, grey line). This model performed consistently well in all tasks. Similarly, the consistently well-performing drive-only model developed drastically different functional specialization between seeds in the auditory tasks (Fig. 6c, blue line). Thus, there are multiple different ways to solve these tasks, and different architectures of feedforward and feedback inputs push the models towards these different solutions.

Another illustrative observation was that the brainlike and reverse models developed more predictable specializations in their visual versus auditory regions compared to other models, i.e. visual regions were largely visual and auditory regions were largely auditory (based on their clustering profiles). This is likely because the brainlike and reverse models had visual-to-auditory connections that were either all feedforward or all feedback, respectively. This further confirms that the distinction between feedforward and feedback inputs, as implemented in our models, helps to determine the set of solutions available to the networks and the regional specializations that they develop.

## 3 Discussion

Neurophysiological and anatomical data suggest that top-down feedback to pyramidal neurons in the neo-cortex has a distinct impact on activity from that of bottom-up feedforward inputs. Specifically, top-down feedback has a more modulatory role, changing the gain of the neurons as well as their spike threshold, rather than directly driving activity (Larkum, 2004; Shai et al., 2015). In this study, we built what is, to our knowledge, the first hierarchical multi-modal deep neural network architecture based on the brain’s anatomy that takes this distinction into account. We then explored how different architectures of feedforward and feedback inputs impacted the behavior of networks on audiovisual integration tasks. We compared a brainlike model, based on human cytoarchitectural data to other models, including a model that reversed the relationship observed in the brain, random models, and a model with only feedforward connections. We found that even in densely connected, identically sized models, different configurations of feedforward and feedback connectivity gave the models different strengths, weaknesses and inductive biases. In particular, deep models with a human brainlike hierarchy exhibited a distinct visual bias, but nevertheless performed well on all audiovisual tasks, qualitatively mimicking a long-known human bias for visual stimuli (Posner et al., 1976; Stokes & Biggs, 2014). While weakly driving composite feedback improved the performance of some of the lagging models, the visual bias of the brainlike model persisted for both composite and multiplicative feedback. Moreover, all regions of the brainlike model developed their expected functional specializations (i.e. visual regions clustered visual stimuli, and auditory regions clustered auditory stimuli), despite us having given no region-specific bias other than connectivity. This suggests that the profile of feedforward and feedback connectivity of a region helps determine its functional specializations. Altogether, our results demonstrate that the distinction between feedforward and feedback inputs has clear computational implications, and that ANN models of the brain should therefore consider top-down feedback as an important biological feature.

### 3.1 Computational impact of top-down feedback

Top-down feedback is speculated to play a variety of roles in the cortex. It’s known to suppress neural responses to predictable inputs (Nassi et al., 2013; Rao & Ballard, 1999)–an observation that forms the core of the predictive coding framework Friston and Kiebel, 2009; Mumford, 1992; Rao and Ballard, 1999. In addition to its predictive function, it’s crucial for modulating attention (Debes & Dragoi, 2023), shaping perception (Manita et al., 2015), and conveying task-specific context (Li et al., 2004; Liu et al., 2021). Longrange feedback from the motor (Jordan & Keller, 2020; Leinweber et al., 2017) and auditory cortex (Garner & Keller, 2021) carry important contextual information to the early visual cortex.

The many proposed roles of top-down feedback have been explored by various computational models (Choksi et al., 2020; Deco & Rolls, 2004; Huang et al., 2020; Jiang & Rao, 2024; Mittal et al., 2020; Pang et al., 2021; Wen et al., 2018), but these previous studies have not generally incorporated the multiplicative, modulatory role of top-down feedback. There are a few important, recent exceptions. First, Naumann et al., 2022 examined the impact of a top-down feedback that modulated the feedforward synapses, showing that modulatory feedback can help to recover source signals from a noisy environment. Second, Wybo et al., 2022 used a composite feedback mechanism and biophysical modeling to show that modulatory feedback can help neurons to flexibly solve multiple linearly inseparable problems through Hebbian plasticity. Next, Islah et al., 2023 showed that multiplicative feedback can provide crucial contextual information to a neural network, allowing it to disambiguate challenging stimuli. Most recently, Tsai et al., 2024 showed that modulatory feedback enhances task-relevant sensory signals in a computational model of S1 and lOFC. Our study adds to this previous work by incorporating modulatory top-down feedback into deep, convolutional, recurrent networks that can be matched to real brain anatomy. Importantly, using this framework we could demonstrate that the specific architecture of top-down feedback in a neural network has important computational implications, endowing networks with different inductive biases.

One other potential computational role for top-down feedback is to provide a credit assignment signal (Greedy et al., 2022; Guerguiev et al., 2017; Lee et al., 2015; Payeur et al., 2021; Roelfsema & Ooyen, 2005; Sacramento et al., 2018). There is some experimental evidence supporting this role for top-down feedback in the brain (Bittner et al., 2017; Doron et al., 2020; Francioni et al., 2023), but it is not yet a widely accepted function for top-down inputs. Nonetheless, if additional experimental evidence supports this role for top-down feedback, it will be critical to also incorporate the credit assignment function into future models.

### 3.2 Testable predictions of audiovisual integration

The idea that auditory cortical regions are higher on the computational hierarchy than visual regions in the human brain is a longstanding, if somewhat controversial hypothesis (King & Nelken, 2009). Neurons in V1 respond strongly to very low-level features, such as oriented gratings (Hubel & Wiesel, 1962). Meanwhile, neurons in A1 respond more strongly to complex, modulated, and relatively long stimuli compared to pure tones and frequency sweeps (Wang et al., 2005). One reason for this may be that the subcortical auditory system in mammals is extensive, and the synaptic distance between the cochlea and A1 is consequently greater than that between the retina and V1. In-line with this, previous modeling work with deep networks showed that the primary auditory area matches intermediate regions of trained deep networks most closely (Kell et al., 2018) while primary visual regions match early layers (Cadena et al., 2019; Khaligh-Razavi & Kriegeskorte, 2014; Yamins & DiCarlo, 2016). Our study suggests that if the hypothesis is true, then there are functional consequences that flow from the architecture of the human brain, most notably, a bias towards relying on visual inputs more readily to resolve ambiguities. This is one clear, experimental prediction that our models make.

However, while humans are generally thought to be a visually-dominant species, some individuals with musical training report a behaviorally and functionally distinct auditory bias (Giard & Peronnet, 1999). The proportion of feedforward and feedback connectivity between the visual and auditory regions may explain these individual differences. This represents another testable prediction flowing from our study, which could be studied in humans by examining the information flow between auditory and visual regions during an audiovisual task (example technique in Pines et al., 2023). If confirmed, the hypothesis provides a concrete, developmentally flexible mechanism for the emergence of human visual bias.

Moreover, the link between connectivity and functional specialization can be studied further within the same framework to produce testable hypotheses about the effects of cortical lesions, implant responses and stroke recovery. Previous large-scale computational models of lesions lend great insight into their effects on brain function, but cannot readily probe their behavioral effects (Alstott et al., 2009; Martínez-Molina et al., 2024). In contrast, the ANNs used in our study can be trained on very complex tasks via backpropagation, and thus could provide direct insight into the potential impact of lesions on task performance and behavior. In general, with intelligent selection of architectures guided by anatomy and tasks selected based on real experimental conditions, ANNs with modulatory top-down feedback and brainlike connectivity could be used to generate many predictions about functional connectivity and lesion recovery.

### 3.3 Limitations and future directions

We made various simplifying assumptions about connectivity in the human brain to build the brain-based model. Feedforward-feedback directionality of a connection is determined in animals using tracer injections, but in human brains, it must be determined using proxy measures. To determine the directionalities in our model, we used externopyramidisation–the relative thickness and differentiation of supragranular layers, known to be highest at the bottom of the sensory hierarchy. While our measure replicates expected hierarchical ordering of brain regions based on previous literature, visual regions exclusively sending feedforward information to auditory regions in the human brain is an extrapolation of available cytoarchitectural data.

While the visual bias of the brainlike model was evident across all scenarios, the reverse model did not display a similar task-agnostic auditory bias (Fig. 2e). This is likely because the auditory dataset in our study is smaller, less variable and thus easier to learn than the visual dataset, causing certain models to shortcut their training and rely excessively on it (Fig. 3b and Fig. 3e). A visual bias helps models ignore this shortcut, whereas an auditory bias makes them more susceptible to it and hampers a model’s ability to ignore audio input on visual tasks (Fig. 2d, Fig. 5b). This points to another limitation in our study, namely the use of relatively simple stimuli and small datasets. Ideally, future work would use rich, natural audiovisual inputs and large datasets to train the networks. This may lead to some different behaviors in the models. Moreover, our results relate to other findings that task demands often supersede architectural choices in computational modeling (Lindsay et al., 2022). Nonetheless, despite these limitations, our model shows clearly that within certain tasks the biases imparted by architectural choices have important implications that interact with the data.

Another important limitation to note is that many aspects of our model are not biologically plausible. Whether a connection is feedback or feedforward is fixed at the start of the experiments, whereas connectivity is more flexible and task-based in biological brains. Moreover, out model uses end-to-end backpropagation and supervised learning. However, our models are not only applicable in these cases - one could easily train the ANNs we designed with different learning algorithms. As well, the models we developed are agnostic to training method and can easily be trained with self-supervised and reinforcement learning tasks. But, we believe that the ability to train our models with backpropagation is important, because that will allow them to learn a wide range of complicated, more natural tasks in future research. We have released the codebase to construct these models to facilitate such research.

Finally, another key limitation in this study is that we did not compare our models directly to human neural data. Our results show clearly that the models’ internal representations are altered by top-down feedback (Fig. 6), so we would expect it to also have an impact on the ability of the models to match the representations in real brains. But, we leave this as future work, which is made easier by the release of the codebase.

In summary, our study shows that modulatory top-down feedback and the architectural diversity enabled by it can have important functional implications for computational models of the brain. We believe that future work examining brain function with deep neural networks should therefore consider incorporating top-down modulatory feedback into model architectures when appropriate. More broadly, our work supports the conclusion that both the cellular neurophysiology and structure of feedback inputs have critical functional implications that need to be considered by computational models of brain function.

## 4 Methods

### 4.1 Model

Each area/layer of the model is a modified Convolutional Gated Recurrent Unit (ConvGRU) cell. A Gated Recurrent Unit is a standard neural network with locally recurrent connections. The processes are identical to that of a typical GRU with the exception of the linear operators, which are replaced by convolutions (represented by the star) as per Ballas et al., 2016, and the topdown signal (m), which modulates the hidden state after it has been reset and combined with the bottom-up input.

The tasks in this paper are non-sequential; the image and the audio stimuli are presented at the same time to V1 and A1 respectively. At each time step, the areas receiving a bottom-up feedforward input (h_l_*_−_*_1_) get activated. The feedforward input is combined with the hidden state memory of the area (h_l_), and modulated by the excitatory topdown signal (m), which is derived from the feedback input (h_l+1_) if it exists. In case there is no feedback input (i.e the higher order areas have not been active yet), m is a matrix of ones. Resets and updates (Equations 1 and 2) are features of the GRU that implement local recurrence. The full equations are presented below.

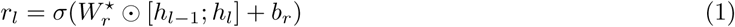

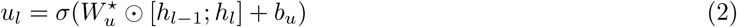

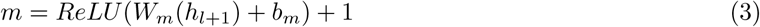

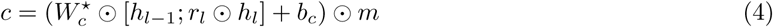

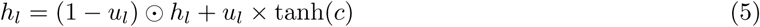

where r_l_ is the reset gate for layer l, u_l_ is the update gate for layer l, h_l_ is the hidden state of layer l, h_l_*_−_*_1_ is the feedforward input from the previous layer, h_l+1_ is the feedback input from the next layer, m is the modulatory top-down signal, c is the candidate activation, [x; y] denotes concatenation along the channel dimension, ★ denotes convolution, ⊙ denotes element-wise multiplication, and W and b represent learnable weights and biases respectively.

The input size of the regions decreased as they moved up the sensory-association hierarchy, i.e V1 and A1 received 32x32 input, while V2 and the auditory belt received 16x16 input. All regions of the model had a hidden state channel size of 10, matching the number of classes. The small size of the model is to make differences in performance clearer for easier tasks, but as shown in Fig. S2, larger models (with hidden state size of 32) have similar image/auditory biases with a higher performance ceiling.

All feedforward inputs to an area were combined and projected into the correct shape by a convolutional projection layer. Feedback inputs were similarly reshaped by a convolutional projection layer to a shape that matches the hidden state of the region.

All models presented in the main figures had the same number of parameters within the two significant figures range (280K), since the number of connections, number of nodes and sizes of hidden states are the same for all of them with only directionality of connections changing. Small differences (¡1K) in size exist between multiplicative and composite feedback models due the extra driving input from high-order areas that are characteristic of the composite models.

#### 4.1.1 Composite feedback

When using composite feedback, the top-down signal was split into the modulatory signal m_mod_matching the shape of the feedforward input, and the driving signal m_d_, which has ten times fewer channels than the feedforward input. In other words, in addition to multiplicative modulation, the feedback signal provided a driving signal at one tenth the strength of the feedforward driving signal. Equation 4 was then adjusted to incorporate the two different types of top-down signals:

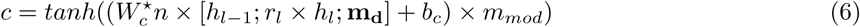

#### 4.1.2 Classification

For supervised classification tasks, the user must designate an output region (IT in visual tasks and A4 in auditory tasks). The hidden states of the output region are fed to two-layer multilayer perceptron (MLP) trained at the same time as the model, which then outputs a classification.

### 4.2 Bases of the human brainlike model

We based the brainlike model on human structural connectivity and histological data. For all human data, regions of interest were determined using the multimodal HCP parcellation (Glasser et al., 2016). Four classic regions in the ventral visual system was chosen to represent the visual cortex: V1, V2, V4, and PIT. Three auditory areas were chosen to represent the auditory cortex: A1, parabelt (part of the belt bordering A4), and A4, a portion of Brodmann area 22 that activated by language tasks and is thought to correspond to area Te3 (Morosan et al., 2005).

To determine the overall connectivity between regions, we used the diffusion tractography data of 50 adult subjects from the Microstructure-Informed Connectomics (MICA-MICs) dataset (Royer et al., 2022). We averaged the connectivity between all pair of regions across subjects and generated a group-average binary connectivity matrix using distance-based thresholding (Betzel et al., 2019).

**Table 1:** Parameters for the modified ConvGRU equations.

| Parameter | Definition |
| --- | --- |
| $r_l$ | Reset gate for layer $l$ |
| $u_l$ | Update gate for layer $l$ |
| $h_l$ | Hidden state of layer $l$ |
| $h_{l-1}$ | Feedforward input from the previous layer |
| $h_{l+1}$ | Feedback input from the next layer |
| $m$ | Modulatory top-down signal |
| $m_{mod}$ | Modulatory component of top-down signal (composite feedback) |
| $m_d$ | Driving component of top-down signal (composite feedback) |
| $c$ | Candidate activation |
| $W_r$ | Learnable weight matrix for reset gate |
| $W_u$ | Learnable weight matrix for update gate |
| $W_c$ | Learnable weight matrix for candidate activation |
| $W_m$ | Learnable weight matrix for modulatory signal |
| $b_r$ | Bias vector for reset gate |
| $b_u$ | Bias vector for update gate |
| $b_c$ | Bias vector for candidate activation |
| $b_m$ | Bias vector for modulatory signal |
| $\sigma$ | Sigmoid activation function |
| $\tanh$ | Hyperbolic tangent activation function |
| ReLU | Rectified Linear Unit activation function |
| $[x; y]$ | Concatenation along the channel dimension |
| $\star$ | Convolution operation |
| $\odot$ | Element-wise multiplication |

To determine the directionality of connectivity between regions, we used the BigBrain quantitative 3D laminar atlas of the human cerebral cortex (Amunts et al., 2013; Wagstyl et al., 2020). We used externopyramidisation (Sanides, 1962), the relative size of supragranular neurons compared to infragranular neurons, as the proxy for the hierachical position of area. In mammals including humans, it is high in regions that send more supragranular feedforward projections and lower in areas that send more infragranular feedback projections (Goulas et al., 2018). This value can be estimated from histological data using the relative laminar thickness and staining intensity of supragranular layers, a process described by Paquola et al., 2020. Similar to the method described in that paper, we rescaled supragranular staining intensity (in) and thickness values to a range of 0 to 1, and used the normalized peak intensity from 20 sample locations (n) and relative thickness of supragranular layers to calculate externopyramidisation.

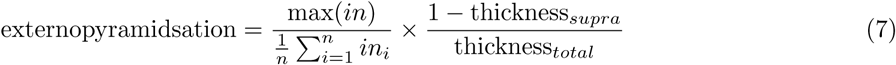

Regions with higher externopyramidisation values were configured to send feedforward connections to and receive feedback from regions with lower externopyramidisation values.

### 4.3 Stimuli generation and preprocessing

We used MNIST handwritten digit database as the unambiguous visual stimuli and the Free Spoken Digit Dataset (FSDD) as the unambiguous auditory stimuli. The visual stimuli were minimally preprocessed, while the audio was Fourier-transformed and converted to log scale to generate uniformly sized mel-spectrograms. The ambiguous visual stimuli were created by Islah et al., 2023 and used with permission. Details of it can be found in the cited paper. The ambiguous visual stimuli all have two equally possible labels, capping the performance of classifiers at around 50 percent if they are not given additional clues.

#### 4.3.1 Ambiguous auditory stimuli

The ambiguous auditory stimuli were generated in a similar manner to the ambiguous visual stimuli. We trained a conditional variational autoencoder (CVAE) (Sohn et al., 2015) to project mel-spectrograms of FSDD data into a 32 dimensional latent space. We then sampled the Euclidean mean of two random data in the latent space with differing labels, and used the decoder of the trained CVAE to deconvolve it into a mel-spectrogram. The resulting ambiguous digit often retained the features of one digit, but lost all features of the other it was “mixed” with. As such, while the ambiguous visual stimuli have two equally possible interpretations by design, the ambiguous auditory stimuli have a single possible interpretation that is nevertheless difficult to identify without additional clues. To ensure a balance of recognizability and ambiguity in the auditory data, we trained a separate softmax classifier on mel spectrograms of unambiguous FSDD data. The softmax classifier output a prediction between 0 and 1.0 (full certainty) for each label. We asked the classifier to predict the labels of each newly generated ambiguous mel spectrogram, and kept only the data and labels that caused the classifier to output a prediction between 0.45 and 0.55. We reiterated the process until we had 4000 unique ambiguous auditory stimuli, the size of the holdout portion of original FSDD dataset. Due to the higher power of the ConvGRU-based models used in our experiments compared to the softmax classifier used for quality control, the models used in the experiments are able to decipher the ambiguous audio up to 80 percent accuracy without clues.

### 4.4 Audiovisual task training

We combined the four types of stimuli into fixed audiovisual datasets, where each datapoint consisted of a pair of audio and visual stimuli and their corresponding labels. We held out 15 percent of each dataset for testing before creating the audiovisual datasets, meaning no image or audio during testing was encountered during training. We calculated the gradient in minibatches of 32, and used the Adam optimizer at learning rate 0.0001 to train the model. All models were trained for 50 epochs. For each experimental condition, we ran 10 different seeds. We additionally tested learning rates of 0.001, 0.01 and 0.1 after getting the primary results. The models show the same behaviors and preferences under all rates except the highest learning rate of 0.1, where all models train poorly.

#### 4.4.1 Training tasks

For tasks VS1 and VS2, we trained the models on a shuffled mix of datasets UAM (unambiguous audio, ambiguous image, matching label) and UUN (unambiguous audio, unambiguous image, mismatched label). We set the label of the image as the target and calculated the cross-entropy loss between it and the model prediction.

For tasks AS1 and AS2, we trained the models on a shuffled mix of datasets AUM (unambiguous audio-ambiguous image, matching label) and UUN (unambiguous audio, unambiguous image, mismatched label). We set the audio labels as the target and otherwise trained the models under the same parameters as the visual tasks.

#### 4.4.2 Control tasks and alignment

Datasets AUN (ambiguous audio, unambiguous image, nonmatching label) and UAN (unambiguous audio, ambiguous image, nonmatching label) were used only for testing in scenarios AS3 and VS3 respectively. The visual alignment of a model is determined as the percentage of time when the model outputs the label of the visual stimuli. Same is true for the auditory dataset. The test-only datasets were also generated solely from holdout stimuli.

In addition, models were separately trained and evaluated on unambiguous MNIST and FSDD data to establish their baseline performance (scenarios VS4 and AS4).

#### 4.4.3 Flexible task training

For the flexible task-switching experiments, the models are given a new output region and trained on datasets AUM, UAM, and UUN. Its output region is given a binary attention flag indicating the input stream to attend to.

After training, the image and auditory alignment of the models were tested using the previously unseen AAN (ambiguous audio, ambiguous image, nonmatching label) dataset.

#### 4.4.4 Process time

All models have a unique hyperparameter called process time, which roughly determines how many computational steps the model can take before it must output an answer. A process time of 1, for instance, will mean that only the primary areas A1 and V1 will have received any feedforward input before the model generates an output. For all audiovisual tasks, we picked a process time equal to the number of areas in the model (7).

### 4.5 Regional activity analysis

We fetched the hidden state representation of all regions of the trained models at 7 different process times from 1-7 while they performed the tasks they were trained on. We then measured the quality of clustering in the latent space of each model region and time point by calculating the Neighborhood Hit (NH) score (Paulovich et al., 2008; Rauber et al., 2017). The NH score is the mean percentage of neighbors r_j_ for each datapoint n who share the same label C[x_i_]. It’s calculated as below, using k=4 for every task.

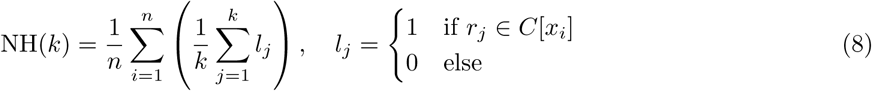

### 4.6 Statistical analysis

Mean final accuracies for multiplicative and composite feedback were compared using Welch’s t-test. Error bars and bands represent standard error of the mean, unless stated otherwise. *, p < 0.05; **, p < 0.01; ***, p < 0.001; ****, p < 0.0001; n.s., not significant.

### 4.7 Code availability

The toolbox used to convert graphs to top-down recurrent neural networks (Connectome-to-Model) is publicly available at https://github.com/masht18/connectome-to-model. The task training scripts and graphs are also available at the same repository.

## 5 Acknowledgements

We thank Nizar Islah for the ambiguous visual dataset as well as discussion regarding the ambiguous auditory dataset. We additionally thank Jessica Royer and Boris Bernhardt for discussions regarding the brain basis of the model, and Colin Bredenberg for helpful comments on the manuscript. This work was supported by NSERC (Discovery Grant: RGPIN-2020-05105; Discovery Accelerator Supplement: RGPAS-2020-00031; Arthur B. McDonald Fellowship: 566355-2022), CIFAR (Canada AI Chair; Learning in Machine and Brains Fellowship), and the Canada First Research Excellence Fund (CFREF Competition 2, 2015-2016) awarded to the Healthy Brains, Healthy Lives initiative at McGill University, through the Helmholtz International BigBrain Analytics and Learning Laboratory (HIBALL). E.B.M. was additionally supported by the Institute for Data Valorization (IVADO), the Centre de recherche Azrieli du CHU Sainte-Justine (CRACHUSJ), Fonds de Recherche du Québec–Santé (FRQS), and CIFAR (Canada AI Chair Mila). This research was enabled in part by support provided by Calcul Québec (https://www.calculquebec.ca/en/) and the Digital Research Alliance of Canada (https://alliancecan.ca/en). The authors acknowledge the material support of NVIDIA in the form of computational resources. M. T received additional support from the Healthy Lives Healthy Brains Graduate Fellowship and UNIQUE Excellence Fellowship.

## 6 Supplemental

**Table 2:** Mean and standard deviation of final epoch accuracy across VS tasks for small models with multiplicative feedback (Fig. 2).

|  | Brainlike | Reverse | Rand1 | Rand2 | Rand3 |
| --- | --- | --- | --- | --- | --- |
| VS1 | $0.868 \pm 0.009$ | $0.914 \pm 0.011$ | $0.885 \pm 0.013$ | $0.880 \pm 0.030$ | $0.836 \pm 0.011$ |
| VS2 | $0.933 \pm 0.006$ | $0.945 \pm 0.013$ | $0.909 \pm 0.021$ | $0.884 \pm 0.058$ | $0.800 \pm 0.052$ |
| VS3 (image align) | $0.446 \pm 0.007$ | $0.440 \pm 0.006$ | $0.436 \pm 0.010$ | $0.424 \pm 0.016$ | $0.406 \pm 0.027$ |
| VS3 (audio align) | $0.216 \pm 0.008$ | $0.242 \pm 0.011$ | $0.237 \pm 0.007$ | $0.247 \pm 0.008$ | $0.255 \pm 0.029$ |
| VS4 (image recognition) | $0.964 \pm 0.003$ | $0.939 \pm 0.007$ | $0.967 \pm 0.002$ | $0.907 \pm 0.025$ | $0.959 \pm 0.009$ |

**Table 3:** Mean and standard deviation of final epoch accuracy across AS tasks for small models with multiplicative feedback (Fig. 3)

|  | Brainlike | Reverse | Rand1 | Rand2 | Rand3 |
| --- | --- | --- | --- | --- | --- |
| AS1 | $0.972 \pm 0.006$ | $0.880 \pm 0.046$ | $0.968 \pm 0.039$ | $0.879 \pm 0.047$ | $0.887 \pm 0.054$ |
| AS2 | $0.944 \pm 0.016$ | $0.937 \pm 0.009$ | $0.945 \pm 0.005$ | $0.945 \pm 0.015$ | $0.946 \pm 0.007$ |
| AS3 (image alignment) | $0.509 \pm 0.066$ | $0.164 \pm 0.102$ | $0.452 \pm 0.181$ | $0.165 \pm 0.095$ | $0.204 \pm 0.137$ |
| AS3 (audio alignment) | $0.326 \pm 0.053$ | $0.775 \pm 0.155$ | $0.428 \pm 0.207$ | $0.764 \pm 0.157$ | $0.729 \pm 0.187$ |
| AS4 (audio recognition) | $0.928 \pm 0.012$ | $0.938 \pm 0.010$ | $0.936 \pm 0.016$ | $0.919 \pm 0.022$ | $0.922 \pm 0.015$ |

**Table 4:** Mean and standard deviation across VS tasks for small models with composite feedback (Fig. 4).

|  | Brainlike | Reverse | Rand1 | Rand2 | Rand3 |
| --- | --- | --- | --- | --- | --- |
| VS1 | $0.928 \pm 0.014$ | $0.907 \pm 0.007$ | $0.946 \pm 0.007$ | $0.945 \pm 0.008$ | $0.941 \pm 0.012$ |
| VS2 | $0.949 \pm 0.010$ | $0.941 \pm 0.012$ | $0.962 \pm 0.007$ | $0.671 \pm 0.402$ | $0.811 \pm 0.317$ |
| VS3 (image align) | $0.442 \pm 0.007$ | $0.445 \pm 0.012$ | $0.450 \pm 0.006$ | $0.329 \pm 0.163$ | $0.384 \pm 0.128$ |
| VS3 (audio align) | $0.247 \pm 0.012$ | $0.235 \pm 0.011$ | $0.247 \pm 0.008$ | $0.483 \pm 0.326$ | $0.368 \pm 0.260$ |
| AS1 | $0.967 \pm 0.007$ | $0.897 \pm 0.054$ | $0.964 \pm 0.038$ | $0.892 \pm 0.053$ | $0.877 \pm 0.059$ |
| AS2 | $0.928 \pm 0.013$ | $0.928 \pm 0.008$ | $0.940 \pm 0.011$ | $0.935 \pm 0.010$ | $0.933 \pm 0.014$ |
| AS3 (image align) | $0.453 \pm 0.073$ | $0.200 \pm 0.112$ | $0.444 \pm 0.137$ | $0.193 \pm 0.128$ | $0.192 \pm 0.130$ |
| AS3 (audio align) | $0.368 \pm 0.065$ | $0.722 \pm 0.167$ | $0.423 \pm 0.164$ | $0.741 \pm 0.173$ | $0.741 \pm 0.189$ |

**Figure S1:**
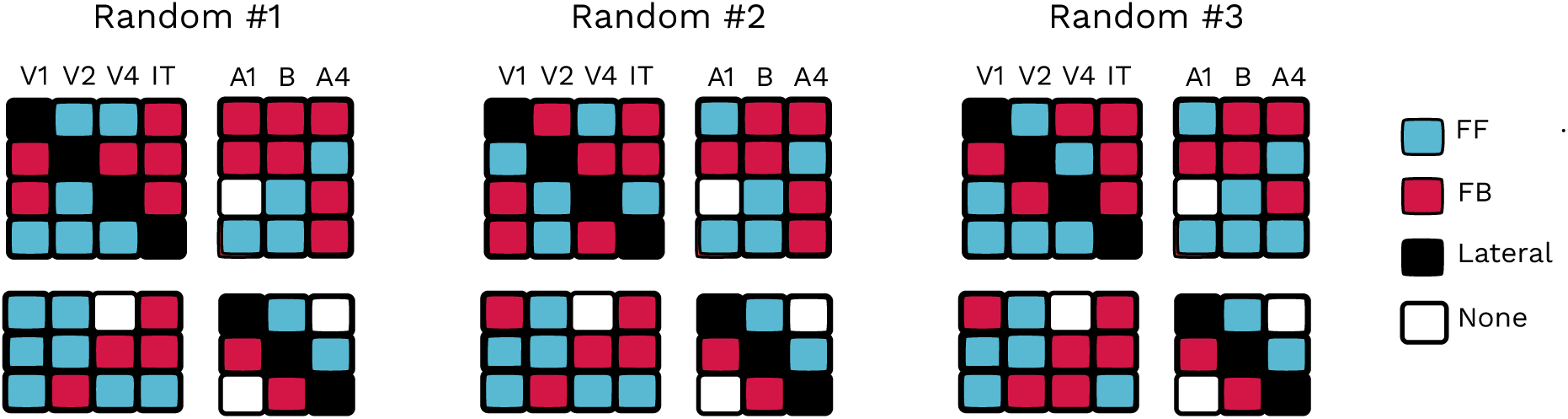
Connectivity of the three random models. The connectivity of the random models were consistent throughout all experiments

**Figure S2:**
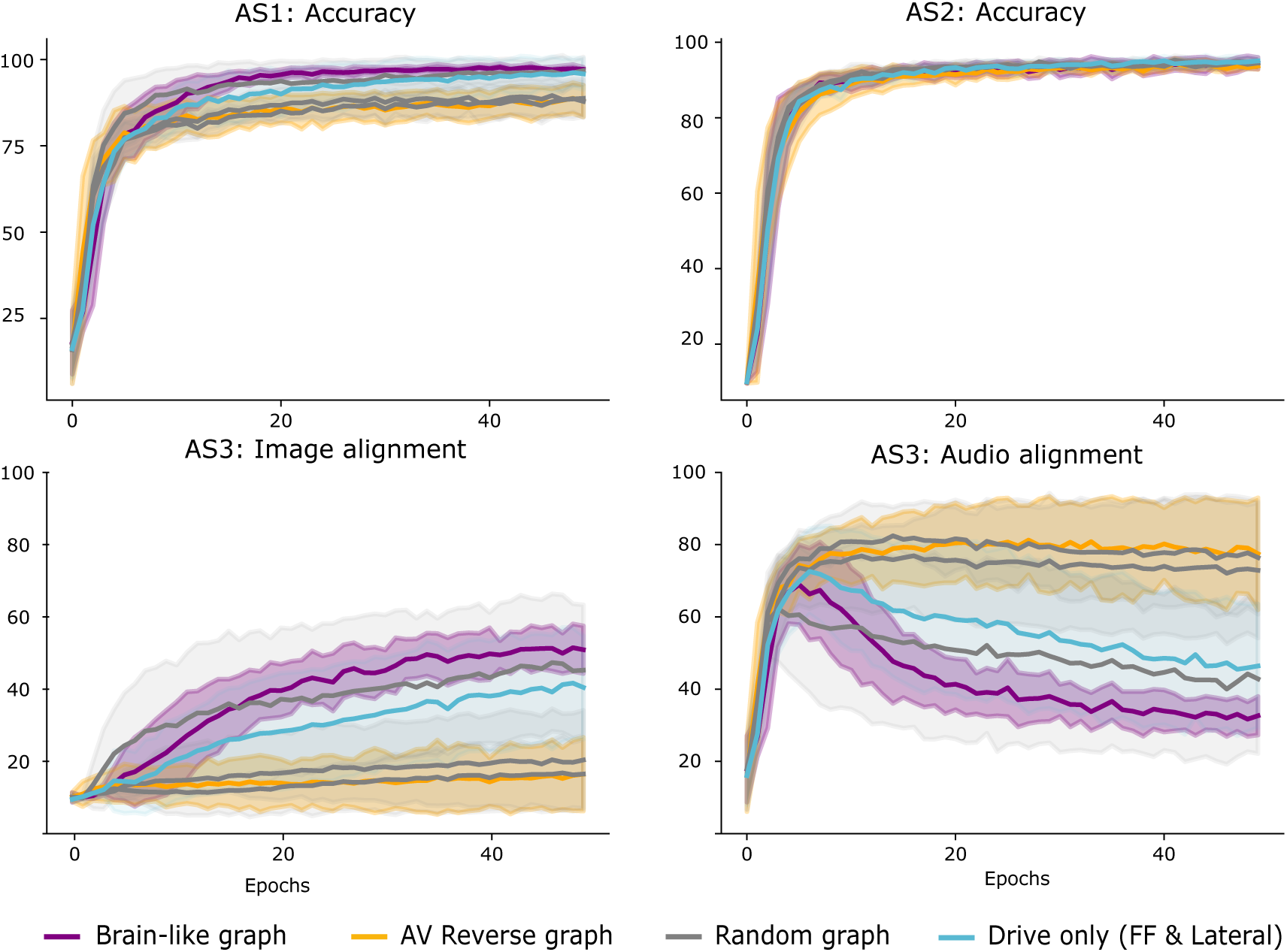
Performance of larger model (hidden state size 32) on AS tasks. Performance is increased across the board, but difference in stimuli preference is still visible

**Figure S3:**
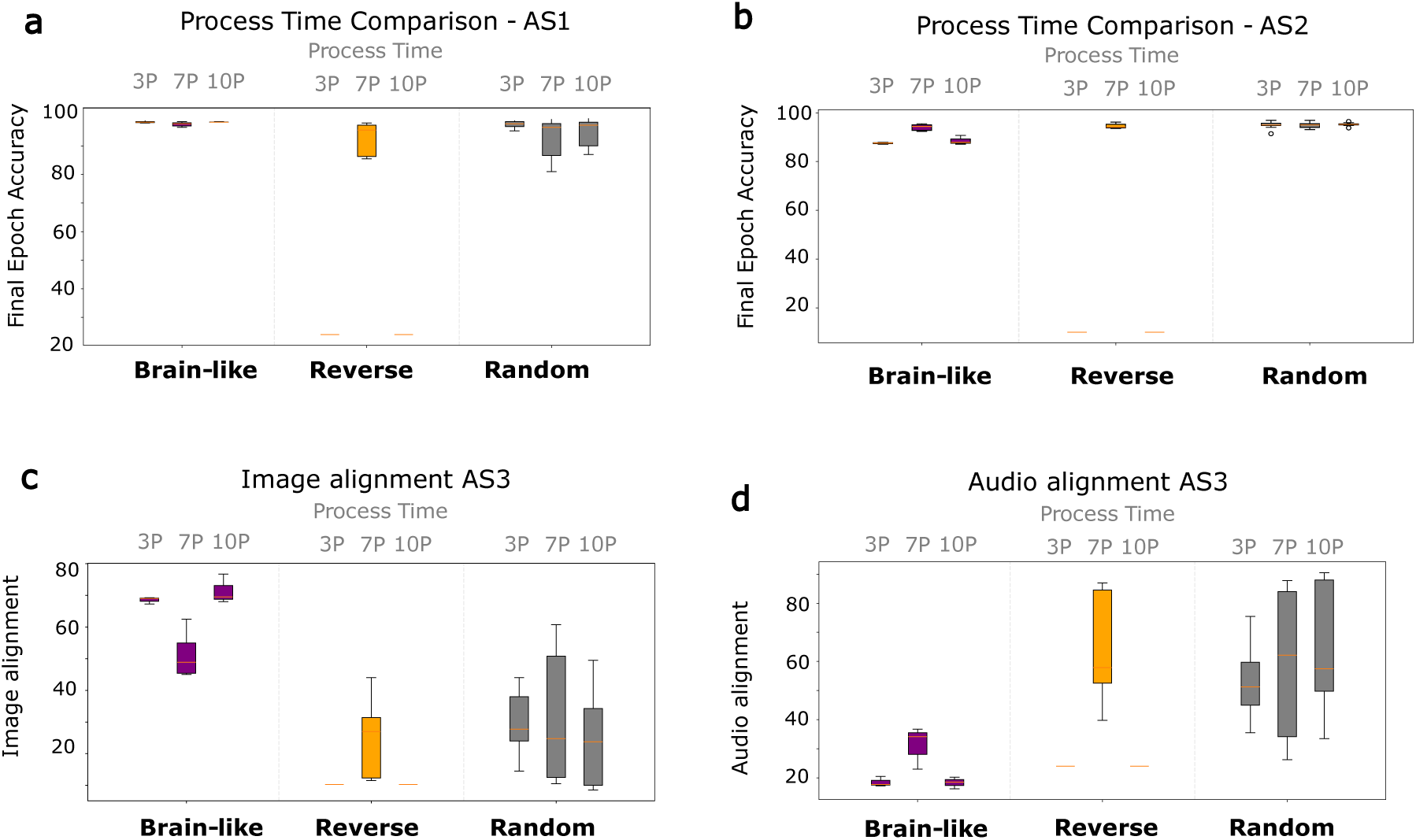
Effect of process time on AS task performance. Final epoch accuracy in model given short (3P), standard (7P, equal to number of model layers) and long (10P) processing times when given **a** ambiguous audio, helpful visual clue, **b** clean audio and distracting image, **c,d** ambiguous audio and nonmatching visual input

**Figure S4:**
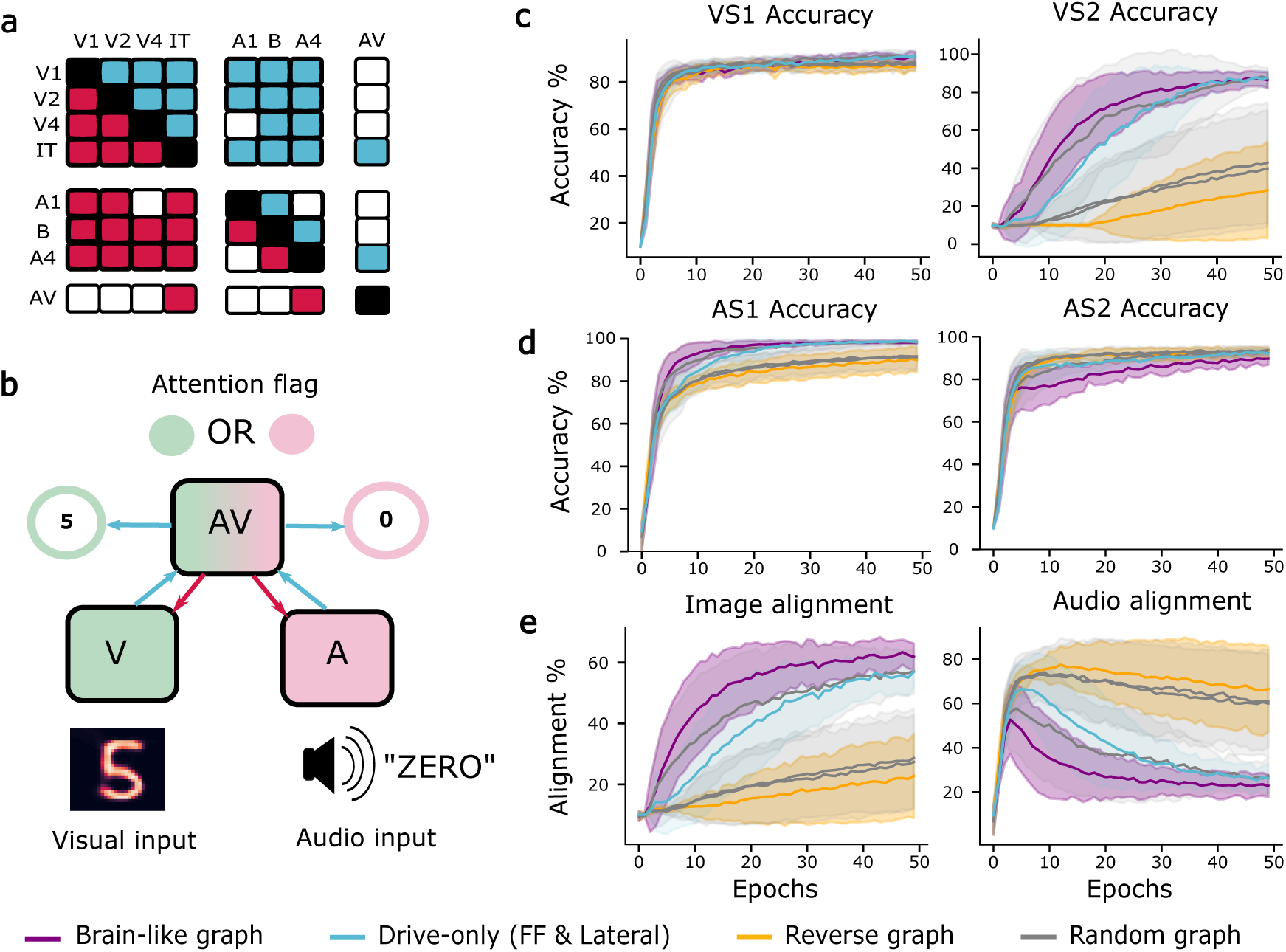
Models with multiplicative feedback on audiovisual switching task. The task and the drive-only (blue) models are the same between this figure and Figure 5.

**Table 5:** Mean and standard deviation of final epoch accuracy across audiovisual switching tasks (Fig. 5)

|  | **Brainlike** | **Reverse** | **Rand1** | **Rand2** | **Rand3** |
| --- | --- | --- | --- | --- | --- |
| VS1 | $0.909 \pm 0.020$ | $0.869 \pm 0.023$ | $0.912 \pm 0.022$ | $0.883 \pm 0.027$ | $0.875 \pm 0.028$ |
| VS2 | $0.864 \pm 0.042$ | $0.285 \pm 0.254$ | $0.874 \pm 0.055$ | $0.399 \pm 0.306$ | $0.430 \pm 0.319$ |
| AS1 | $0.982 \pm 0.008$ | $0.900 \pm 0.058$ | $0.983 \pm 0.014$ | $0.917 \pm 0.062$ | $0.913 \pm 0.065$ |
| AS2 | $0.897 \pm 0.029$ | $0.925 \pm 0.023$ | $0.922 \pm 0.018$ | $0.937 \pm 0.017$ | $0.937 \pm 0.019$ |
| Image align | $0.619 \pm 0.044$ | $0.228 \pm 0.138$ | $0.570 \pm 0.073$ | $0.273 \pm 0.155$ | $0.286 \pm 0.163$ |
| Audio align | $0.228 \pm 0.050$ | $0.665 \pm 0.193$ | $0.273 \pm 0.083$ | $0.611 \pm 0.206$ | $0.600 \pm 0.242$ |

**Table 6:** Mean and standard deviation of final epoch accuracy for big models on AS tasks (Fig. S2)

|  | **Brainlike** | **Reverse** | **Rand1** | **Rand2** | **Rand3** |
| --- | --- | --- | --- | --- | --- |
| AS1 | $0.987 \pm 0.004$ | $0.949 \pm 0.036$ | $0.984 \pm 0.008$ | $0.979 \pm 0.008$ | $0.977 \pm 0.010$ |
| AS2 | $0.952 \pm 0.011$ | $0.949 \pm 0.009$ | $0.953 \pm 0.011$ | $0.962 \pm 0.012$ | $0.971 \pm 0.002$ |
| AS3 (image align) | $0.556 \pm 0.036$ | $0.201 \pm 0.128$ | $0.358 \pm 0.102$ | $0.366 \pm 0.045$ | $0.353 \pm 0.105$ |
| AS3 (audio align) | $0.280 \pm 0.015$ | $0.689 \pm 0.203$ | $0.477 \pm 0.124$ | $0.431 \pm 0.045$ | $0.484 \pm 0.135$ |

